# Leukemia cell of origin influences apoptotic priming and sensitivity to LSD1 inhibition

**DOI:** 10.1101/2020.05.11.089060

**Authors:** Sheng F. Cai, S. Haihua Chu, Aaron D. Goldberg, Salma Parvin, Richard P. Koche, Jacob L. Glass, Eytan M. Stein, Martin S. Tallman, Filiz Sen, Christopher Famulare, Monica Cusan, Chun-Hao Huang, Chun-Wei Chen, Lihua Zou, Keith B. Cordner, Nicole L. DelGaudio, Vidushi Durani, Mitali Kini, Madison Rex, Helen S. Tian, Johannes Zuber, Timour Baslan, Scott W. Lowe, Hugh Y. Rienhoff, Anthony Letai, Ross L. Levine, Scott A. Armstrong

## Abstract

Previous studies have established that the cell of origin of oncogenic transformation is a determinant of therapeutic sensitivity. However, the mechanisms governing cell-of-origin-driven differences in therapeutic response have not been delineated. Leukemias initiating in hematopoietic stem cells (HSC) are less sensitive to cytotoxic chemotherapy and express high levels of the transcription factor *Evi1* compared to leukemias derived from myeloid progenitors. Here, we compared drug sensitivity and expression profiles of murine and human leukemias initiated in either HSCs or myeloid progenitors to reveal a novel function for *Evi1* in modulating p53 protein stability and activity. HSC-derived leukemias exhibit decreased apoptotic priming, attenuated p53 transcriptional output, and resistance to lysine-specific demethylase 1 inhibitors in addition to classical genotoxic stresses. p53 loss-of-function in *Evi1*^low^ progenitor-derived leukemias induces resistance to LSD1 inhibition. By contrast, *Evi1*^high^ leukemias are sensitized to LSD1 inhibition by the BH3 mimetic venetoclax, resulting in enhanced apoptosis and greater reductions in disease burden. Our findings demonstrate a cell-of-origin determined novel role for *EVI1* in p53 wild-type cancers in reducing p53 function and provide a strategy to circumvent drug resistance in high-risk, chemoresistant *EVI1*^high^ AML.

---

Cancers arise through the acquisition of genetic alterations that transform normal cells into malignant clones. While the genetic drivers of cancer have been extensively described, the role of the cell of origin in malignant transformation is less well characterized. Previous studies established that epigenetic states conferred by cell of origin shape mutational profiles^1^ and molecular classification across a diverse array of cancer subtypes^2^. Mouse models of acute myeloid leukemia (AML) provided evidence that manipulation of cell of origin through transformation of specific hematopoietic compartments can give rise to leukemias with distinct phenotypic features, though similar studies validating this in human AML with different cells of origin have not yet been reported. AML derived from hematopoietic stem cells (HSC), marked by high expression of the oncogenic transcription factor *Evi1*, exhibit higher disease penetrance, aggressiveness, and resistance to cytotoxic chemotherapy when compared to leukemias arising from more differentiated hematopoietic progenitor cells^3-6^. These findings correlate with clinical outcomes, as *EVI1* overexpression was the single prognostic factor associated with inferior overall survival among AML patients harboring *MLL* gene rearrangements^7,8^.

Here, we report that the cell of origin of leukemia initiation influences p53 activity and, in turn, apoptotic priming in both mouse and human models of AML. This differential p53 activity is modulated at the level of protein abundance, with greater p53 protein expression in AML arising from granulocyte-monocyte progenitor (GMP) cells when compared to HSC-derived leukemias. We demonstrate that cell-of-origin-dependent p53 protein expression influences the differential sensitivity of these leukemias to genotoxic stresses including chemotherapy^3-5^ and ionizing radiation, as well as to targeted epigenetic therapies that inhibit lysine-specific demethylase 1 (LSD1). LSD1 is a histone demethylase implicated in DNA damage responses and the p53 pathway^9,10^ and is a therapeutic target in AML for which pharmacologic inhibitors are currently being investigated in early-phase clinical trials^11-14^. We also show that resistant *Evi1*^high^ HSC-derived leukemias can be further sensitized by combining an LSD1 inhibitor with the BH3 mimetic venetoclax. Taken together, our findings comparing leukemias arising from different cells of origin led us to discover that the transcription factor *Evi1* can modulate p53 activity in p53 wild-type cancer cells to modulate drug sensitivity, and we identified a therapeutic combination that can elicit durable clinical responses in chemoresistant high-risk *EVI1*-rearranged AML.

## Results

### HSC- and GMP-derived leukemias exhibit differential sensitivity to LSD1 inhibition

To investigate whether cell of origin of leukemic transformation influences sensitivity to LSD1 inhibition, MLL-AF9 was expressed in murine lineage^-^Sca1^+^Kit^+^ (LSK) HSCs or GMPs followed by transplantation into recipient mice. GFP^+^ leukemic blasts were harvested from bone marrow of moribund recipient mice and grown in culture. Consistent with previously published reports^3-5^, LSK-derived murine MLL-AF9 (LSK-MLL-AF9) leukemias were more resistant to doxorubicin exposure when compared to GMP-derived (GMP-MLL-AF9) leukemias (**Supp. Figure 1a**). The leukemias arising from different cells of origin exhibited identical baseline steady-state growth kinetics (**Supp. Figure 1b**), doubling times (**Supp. Figure 1c**), and similar levels of MLL-AF9 mRNA expression (**Supp. Figure 1h**). However, treatment with the irreversible LSD1 inhibitor IMG-7289 revealed marked differences in sensitivity between LSK- and GMP-MLL-AF9 with separation by at least 2 logs in IC_50_ values (**Figure 1a, Supp. Figure 1d**). This differential sensitivity to IMG-7289 was not abrogated by extended treatment duration for as long as 15 days (**Supp. Figure 1d**). We observed similar differential sensitivity to an additional irreversible LSD1 inhibitor, IMG-98 (**Supp. Figure 1e**), suggesting that this phenotype is not attributable to off-target effects. By contrast, pharmacologic inhibition of the H3K79 methyltransferase DOT1L, a known therapeutic vulnerability in MLL-rearranged AML, did not reveal differential sensitivity based on cell of origin (**Supp. Figure 1f**). To test if differential sensitivity to LSD1 inhibition is observed *in vivo*, we transplanted primary LSK- and GMP-MLL-AF9 leukemias into secondary recipients and treated mice via oral gavage with either vehicle control or IMG-7289 (25 mg/kg) for a 14-day period, after which the mice were monitored for survival. While recipient mice engrafted with GMP-MLL-AF9 leukemias exhibited a treatment effect with prolongation of median survival in IMG-7289-treated mice (26 days control vs 33.5 days IMG-7289, p=0.0007, **Figure 1b**), there was no prolongation of median survival in recipient mice transplanted with LSK-MLL-AF9 leukemias (**Figure 1c**). These findings suggest that cell of origin of leukemic transformation affects sensitivity to LSD1 inhibition *in vivo* and *in vitro*.

**Figure 1.**
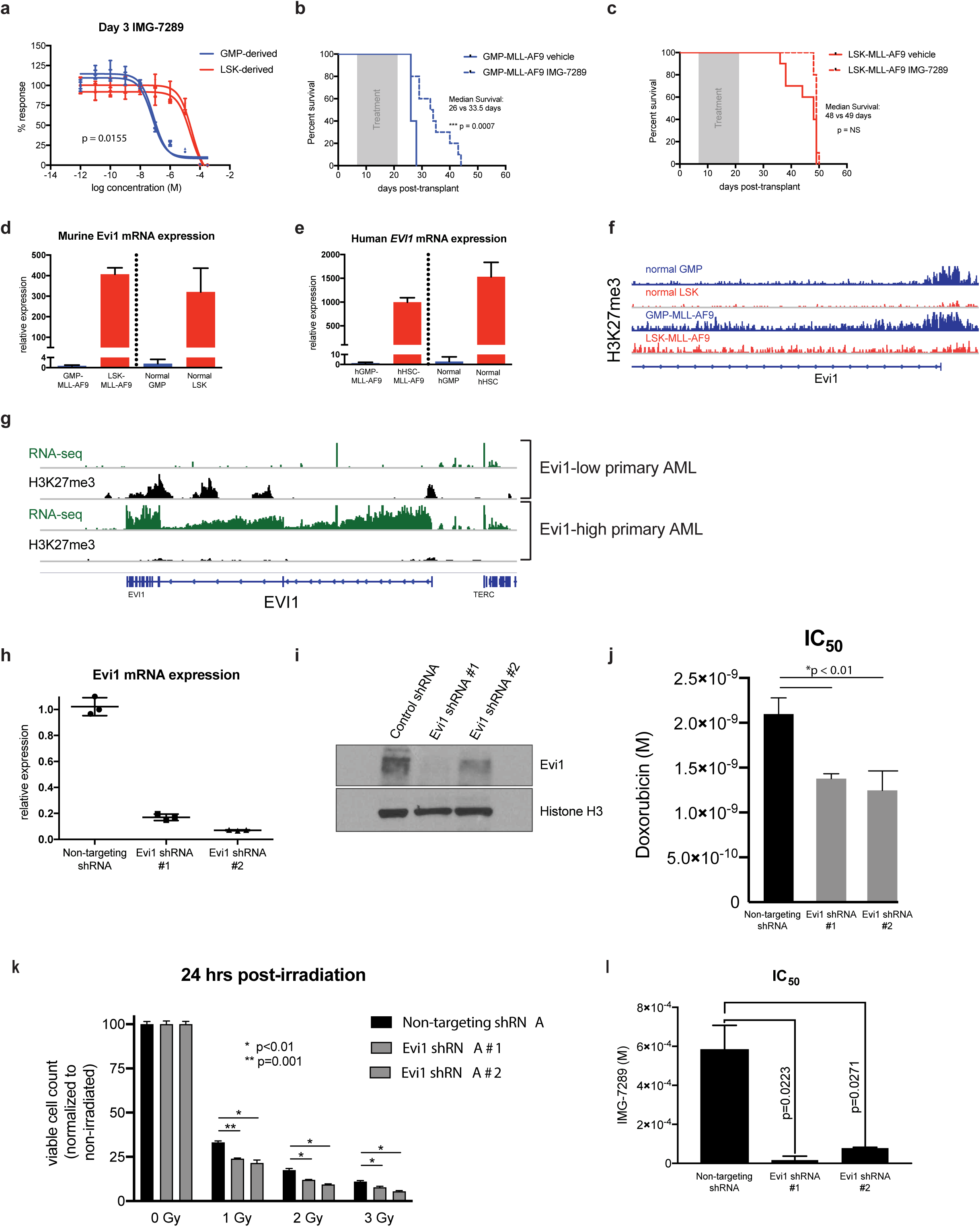
Differential sensitivity of LSK- and GMP-derived MLL-AF9 leukemias to LSD1 inhibition is Evi1-dependent. (**A**) Independent murine LSK- and GMP-derived MLL-AF9 leukemias were treated for 3 days with variable concentrations of IMG-7289 *in vitro*. Cell Titer Glo assays were used to quantitate viable cells cultured in triplicate wells and percent response was derived by normalizing to corresponding cells cultured in DMSO vehicle control. (**B**) Wild-type C57BL/6 mice engrafted with 5×10^4^ syngeneic GMP-MLL-AF9 cells were treated via oral gavage daily with IMG-7289 at a dose of 25 mg/kg (N=12 mice) or vehicle control (N=12 mice) during a 14-day treatment window (grey box). Kaplan-Meier survival curves comparing IMG-7289-treated and vehicle treated mice are shown. (**C**) Wild-type C57BL/6 mice engrafted with 5×10^4^ syngeneic LSK-MLL-AF9 cells were treated via oral gavage daily with IMG-7289 at a dose of 25 mg/kg (N=12 mice) or vehicle control (N=12 mice) during a 14-day treatment window (grey box). Kaplan-Meier survival curves comparing IMG-7289-treated and vehicle treated mice are shown. (**D**) qRT-PCR was performed to measure abundance of Evi1 transcripts in GMP-MLL-AF9, LSK-MLL-AF9, normal sorted GMP cells, and normal sorted LSK cells from C57BL/6 wild-type mice. Bar graphs shown represent pooled biological replicates from 3 independent leukemias and 3 independent sorted LSKs and GMPs. (**E**) qRT-PCR data assessing Evi1 mRNA levels in human cord blood GMP- and HSC-derived MLL-AF9 leukemias engrafted in sublethally irradiated NSG mice and normal human cord blood GMP and HSC cells. (**F**) H3K27me3 ChIP-seq tracks at the Evi1 locus of normal GMP and LSK cells from previously published datasets^21^. Representative H3K27me3 ChIP-seq tracks are also shown from GMP-MLL-AF9 and LSK-MLL-AF9 cells. (**G**) RNA-seq (green) and H3K27me3 (black) ChIP-seq tracks at the *EVI1* locus from *MLL*-rearranged AML patient samples (N=3) exhibiting either high or low expression of *EVI1*. (**H**) qRT-PCR expression analysis of Evi1 from LSK-MLL-AF9 cells transduced with either a non-targeting control shRNA or two independent shRNAs targeting Evi1. (**I**) Western blot analysis of LSK-MLL-AF9 cells expressing control shRNA or Evi1 shRNAs as in H. (**J**) Bar plots representing IC_50_ values from 3 independent Cell Titer Glo doxorubicin kill curve assays using LSK-MLL-AF9 cells expressing non-targeting control shRNA or Evi1 shRNAs. (**K**) Bar plots depicting normalized viable cell counts 24 hours after exposure to ionizing radiation (0, 1, 2, or 3 Gy), comparing LSK-MLL-AF9 cells transduced with either a non-targeting control shRNA or two independent shRNAs targeting Evi1. Data shown are representative of 3 independent experiments. (**L**) Bar plots representing IC_50_ values from 3 independent Cell Titer Glo IMG-7289 kill curve assays using LSK-MLL-AF9 cells expressing non-targeting control shRNA or Evi1 shRNAs.

LSK- and GMP-MLL-AF9 leukemias can be discerned by differential expression of the oncogenic transcription factor *Evi1*, with LSK-MLL-AF9 leukemias having significantly higher *Evi1* mRNA abundance relative to GMP-MLL-AF9 leukemias^3-6^ (**Figure 1d**). Differential *Evi1* expression is also observed in the normal LSK and GMP compartments^15^. Kataoka et al. generated a mouse model in which GFP is knocked into the endogenous locus to allow for the tracking of *Evi1* expression in different hematopoietic compartments^15^. *Evi1* is most highly expressed in long-term HSCs (LT-HSCs), and mice heterozygous for *Evi1* had reduced LT-HSC self-renewal which can be rescued by restoration of *Evi1* expression. Importantly, this GFP reporter mouse showed that Evi1 expression was highly restricted to the LSK fraction with approximately half of the cells in this population expressing GFP. In contrast, virtually no GFP reporter activity was detectable in the GMP and MEP compartments. *EVI1* expression level in AML patients harboring *MLL* gene rearrangements has prognostic value, such that *EVI1*^high^ *MLL*-rearranged AML patients had inferior overall survival, relapse-free survival, and event-free survival^7^. These data are consistent with the hypothesis that prognostic subtypes of human AML can be modeled based on different cells of origin, however this has not been investigated in a human AML model system. To that end, we employed validated human immunophenotypes^16-18^ to sort-purify CD34-selected populations of CD34^+^CD38^-^ enriched human HSCs (hHSC) and CD34^+^CD38^+^CD110^−^CD45RA^+^ human GMP (hGMP) from umbilical cord blood. These populations were then transduced with the MLL-AF9 oncogene and transplanted into sublethally irradiated NOD/SCID/IL-2Rg_c_-null (NSG) mice to generate primary human cord blood-derived MLL-AF9 leukemias. Moribund mice engrafted with high-burden (uniformly >98%) CD33^+^GFP^+^ myeloid blasts in the bone marrow (**Supp. Figure 1g)**, regardless of cell of origin (hGMP or hHSC). Similar to murine MLL-AF9 leukemias, *EVI1* mRNA abundance in hHSC-derived leukemias was higher than *EVI1* transcript levels in hGMP-derived leukemias, and this expression pattern in the leukemias is seen in the corresponding normal human hematopoietic compartments (**Figure 1e**).

Given that differential *Evi1* expression in MLL-AF9 leukemias arising from enriched HSC and GMP populations is observed in the non-malignant stem/progenitor compartments prior to retroviral transduction, we hypothesized that *Evi1* expression levels would correlate with an activated or repressive epigenetic state of the cell of origin prior to oncogenic transformation. The importance of trimethylation of lysine 27 on histone H3 (H3K27me3) as a repressive chromatin mark in regulating gene expression changes during hematopoietic differentiation was previously demonstrated via genetic approaches^19^ and genome-wide studies of histone modifications^20^. We performed H3K27me3 chromatin immunoprecipitation with high-throughput sequencing (ChIP-seq) analysis of three independent murine LSK-derived MLL-AF9 leukemias and three independent GMP-derived leukemias. We also resourced publicly available H3K27me3 ChIP-seq data from normal murine LSK and GMP populations^21^. Concordant with the expression data, the transcriptional start site (TSS) of *Evi1* in both normal mouse GMPs and GMP-derived leukemias were marked by H3K27me3 (**Figure 1f)**. In contrast, mouse LSKs and LSK-derived MLL-AF9 leukemias did not show H3K27me3 enrichment at the TSS of the *Evi1* locus, suggesting that *Evi1* is Polycomb-repressed in GMPs and GMP-MLL-AF9 leukemias but not in LSKs or LSK-MLL-AF9 leukemias. To determine if differential occupancy of the *EVI1* locus by H3K27 trimethylation is also observed in human AML, we performed H3K27me3 ChIP-seq and RNA-seq on primary *MLL*-rearranged AML patient samples. *Evi1*^high^ *MLL*-rearranged AML samples were not marked by H3K27me3 throughout the *EVI1* gene locus, whereas *Evi1*^low^ patient samples were characterized by H3K27 trimethylation (**Figure 1g)**.

Previous studies^6,22^ have shown that knockdown of Evi1 in both mouse and human leukemia cells enhanced sensitivity to cytotoxic chemotherapy and other pro-apoptotic agents. To determine if perturbation of *Evi1* expression in LSK-MLL-AF9 leukemia cells is able to sensitize these cells to LSD1 inhibition, we attenuated *Evi1* expression in LSK-derived AML cells. Two independent shRNA constructs targeting the *Evi1* locus achieved knockdown of greater than 80% relative to non-targeting control shRNAs (**Figure 1h-i**). Consistent with prior reports, *Evi1* knockdown resulted in sensitization to cytotoxic stressors and pro-apoptotic stimuli, such as doxorubicin (**Figure 1j**), ionizing radiation (**Figure 1k**), and the BH3 mimetic venetoclax (**Supp. Figure 1i**). We found that *Evi1* knockdown using these independent shRNA constructs significantly reduced the IMG-7289 IC_50_ by 34-fold and 7.5-fold for *Evi1* shRNAs #1 and #2, respectively (**Figure 1l**). These findings show that the relative sensitivity to LSD1 inhibition observed between LSK- and GMP-derived MLL-AF9 murine leukemias is conferred in part by differential expression of *Evi1*.

### HSC- and GMP-derived leukemias exhibit differential apoptotic priming

To understand the physiological basis for how GMP-MLL-AF9 leukemias are more sensitive to LSD1 inhibition than LSK-MLL-AF9 leukemias, we treated these cells with a fixed concentration of IMG-7289 (1 μM), where the greatest sensitivity threshold between GMP-MLL-AF9 and LSK-MLL-AF9 leukemias was observed (**Figure 1a**), which resulted in a reduction in viable cell counts for GMP-MLL-AF9 cells but not LSK-MLL-AF9 cells (**Figure 2a**). GMP-derived leukemia cells exhibited greater Annexin V staining after IMG-7289 treatment, whereas no such increase in apoptotic cells was observed in LSK-MLL-AF9 cells treated with IMG-7289 (**Figure 2b**). During apoptotic cell death, loss of mitochondrial transmembrane potential (Ψ) occurs before typical morphologic changes associated with apoptosis are observed and also before translocation of phosphatidylserine to the external portion of the cell membrane^23,24^, and this loss of potential can be measured by loss of fluorescence in tetramethylrhodamine ethyl ester (TMRE) stained cells. Loss of TMRE fluorescence was seen in GMP-MLL-AF9 cells following treatment with 1 μM IMG-7289 for 48 hours, represented as an increased proportion of cells with Ψ^low^ TMRE staining (**Figure 2c**). This phenotype was completely blunted in treated LSK-MLL-AF9 cells, with summary plots depicting the proportion of Ψ^low^ cells in **Figure 2d**. The pro-apoptotic effects on MLL-AF9 leukemia cells observed with this class of tranylcypromine-derived LSD1 inhibitors has previously been shown by Harris et al. to closely phenocopy LSD1 knockdown by shRNA^12^.

**Figure 2.**
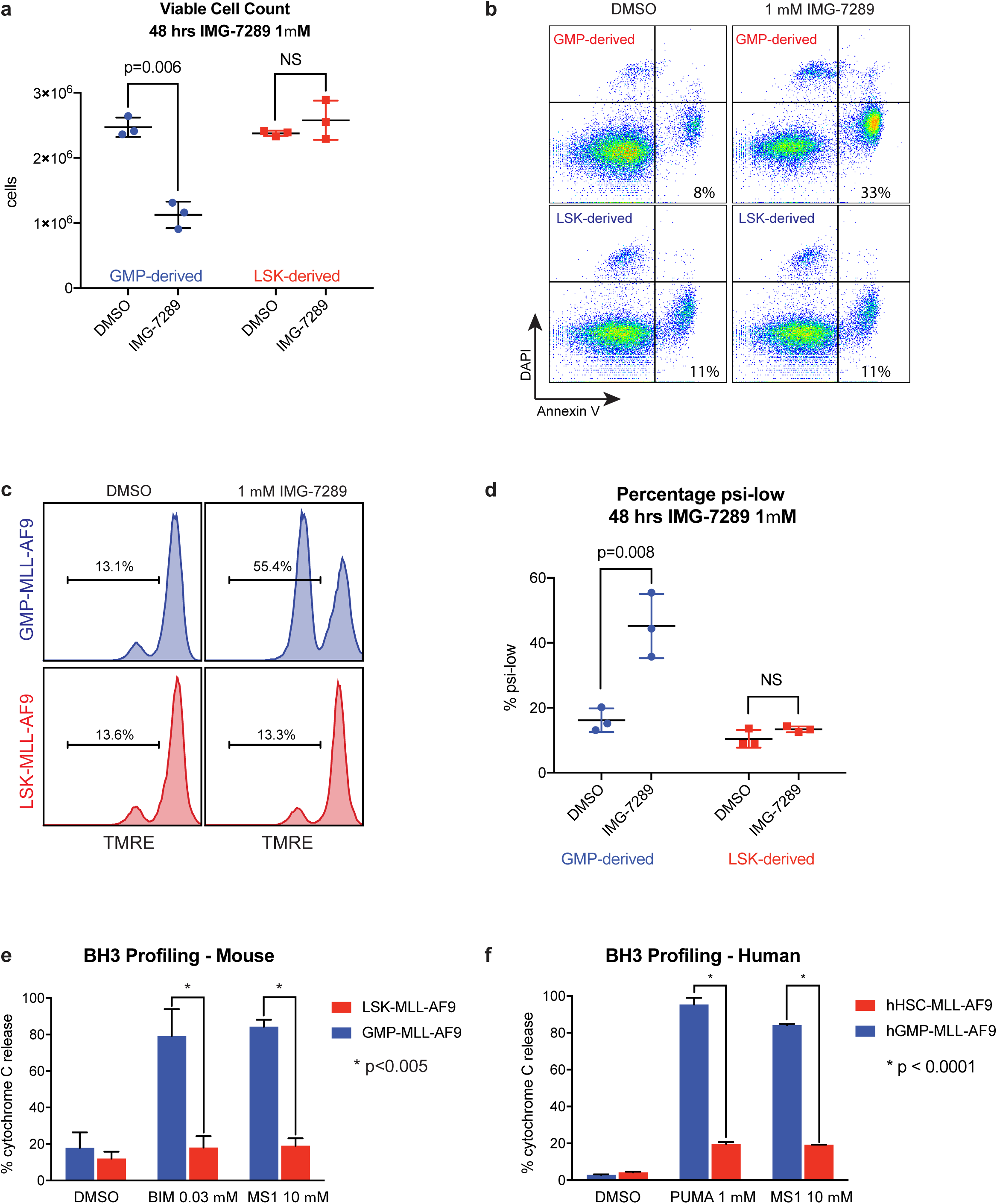
Differential apoptotic thresholds in murine and human MLL-AF9 leukemias arising from distinct cells of origin. LSK- and GMP-derived MLL-AF9 leukemias were treated with DMSO or 1 μM IMG-7289. (**A**) Viable DAPI^-^ cells were enumerated after 48 hours of treatment. (**B**) Annexin V and DAPI staining was assessed by flow cytometry. p<0.03, using Student’s t-test comparing percent Annexin V-positive cells between GMP- and LSK-derived. Data shown are representative of 3 independent experiments. (**C**) TMRE staining was measured by flow cytometry as a readout of mitochondrial membrane potential after 48 hours of treatment. Statistical analyses summarized in (**D**). Data shown are representative of 3 independent experiments. (**D**) Summary scatter plots depicting the percentage of cells within the Ψ^low^ gate defined by TMRE flow cytometry in (**C**). BH3 profiling of (**E**) murine LSK- and GMP-derived MLL-AF9 leukemia cells or (**F**) human cord blood HSC- and GMP-derived leukemias exposed to DMSO control or pro-apoptotic peptides.

In order to functionally interrogate apoptotic priming or triggering thresholds in our MLL-AF9 leukemias derived from murine and human HSCs or GMPs, we employed BH3 profiling, an experimental approach that probes the ability of unique peptides derived from BH3 domains of pro-apoptotic BH3-only protein family members to induce cytochrome *c* release as a readout of apoptosis induction in cells^25,26^. BH3 profiling demonstrates that apoptotic priming increases with myeloid differentiation^27^. The added utility of this assay is that it allows for primary human umbilical cord blood hHSC- and hGMP-derived MLL-AF9 leukemias, which cannot be passaged *in vitro*, to be functionally assessed for apoptotic priming. Murine GMP-derived MLL-AF9 leukemia cells exhibited a high degree of apoptotic priming as assessed by multiple pro-apoptotic peptides (e.g. BIM and MS1) and LSK-derived MLL-AF9 leukemias are more blunted in their ability to trigger apoptosis in response to these same peptides (**Figure 2e, Supp. Figure 2a**). Similarly, human cord blood GMP-derived MLL-AF9 primary leukemias exhibited a higher degree of priming across multiple peptides (e.g. PUMA and MS1), whereas hHSC-derived MLL-AF9 leukemias were more resistant to apoptosis induction (**Figure 2f, Supp. Figure 2b**).

To further explore the link between apoptotic priming and drug sensitivity, we hypothesized that perturbation of apoptotic priming via overexpression of Bcl-2 (**Supp. Figure 3a**) in the more sensitive GMP-MLL-AF9 cells would induce resistance to LSD1 inhibition. GMP-MLL-AF9 cells overexpressing anti-apoptotic Bcl-2 exhibited blunted apoptotic response to a variety of pro-apoptotic peptides relative to cell expressing empty vector control (**Supp. Figure 3b**). Bcl-2-overexpressing cells also had higher IC_50_ upon treatment with IMG-7289 in growth inhibition assays, but also exhibited greater resistance to suppression of viable cell growth by doxorubicin. Together, these data suggest that apoptotic priming underlies the differential sensitivity to LSD1 inhibition and to chemotherapy seen in LSK- and GMP-derived MLL-AF9 leukemias.

### Cell of origin of leukemic transformation influences p53 protein stability in an Evi1-dependent manner

We next investigated the mechanisms that govern cell of origin-specific sensitivity to LSD1 inhibition and differential apoptotic priming. We first conducted a chromatin regulator-focused shRNA screen in murine LSK-derived MLL-AF9 leukemia cells. A library comprised of 2,252 shRNAs targeting 468 known chromatin regulators was constructed in a TRMPV-Neo backbone and transduced as a single pool into Tet-on-competent, monoclonal mouse LSK-derived MLL-AF9 leukemia cells. After neomycin (G418) selection, shRNA expression was induced upon treatment with doxycycline. shRNA-expressing cells were then sorted, and changes in shRNA library representation after 6 days of treatment with IMG-7289 were assessed by high-throughput sequencing of shRNA guide strands as previously described^28^. Mean log_2_-fold change in shRNA representation comparing IMG-7289-treated vs day 0 untreated cells was plotted in rank order, revealing both depleted and enriched shRNAs in IMG-7289-treated cells (**Supp. Table 1**). Among the shRNAs most highly enriched after IMG-7289 treatment was an shRNA targeting p53, suggesting that p53 knockdown can mediate resistance to LSD1 inhibition (**Figure 3a**). Additional shRNA hits that were enriched with LSD1 inhibitor treatment included p53 pathway components *Gadd45a, Csnk2a, Smyd2*, and *Bmi1* (**Supp. Table 1**). *Gadd45a* is a well-known direct transcriptional target of p53 that mediates cell cycle arrest and apoptosis, and Csnk2a1 has also been shown to be required for p53-mediated apoptosis^29^. In contrast, two independent shRNAs targeting the the transcriptional regulator *Cdk9* were depleted in the IMG-7289-treated cells relative to DMSO with log_2_ fold change of −0.914 and - 0.4105, suggesting that loss of *Cdk9* enhances sensitivity to LSD1 inhibition. Other groups have previously demonstrated that attenuation of Cdk9 in MLL-AF9 cells induces apoptosis^30^.

**Figure 3.**
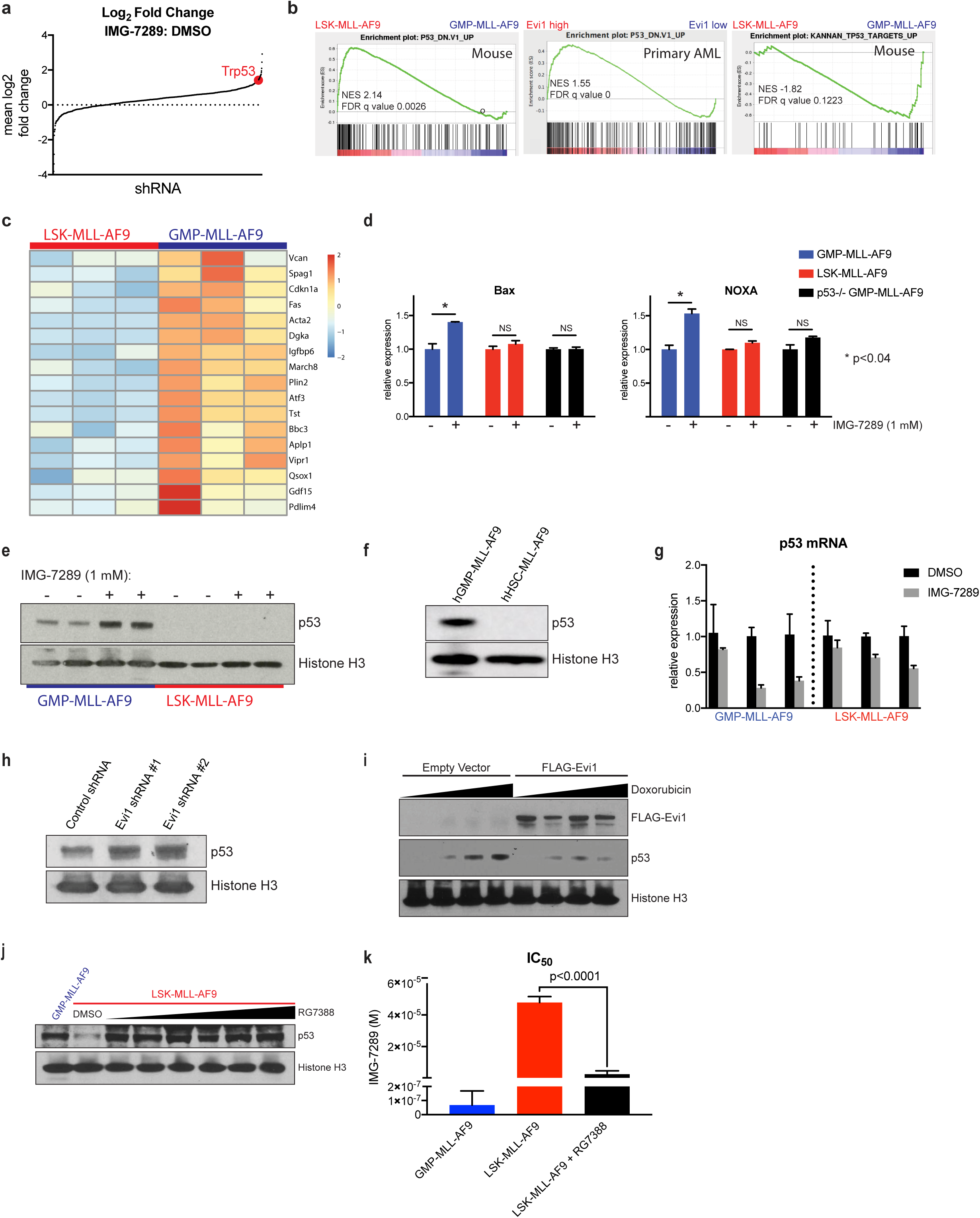
p53 protein abundance and transcriptional output is greater in GMP-MLL-AF9 relative to LSK-MLL-AF9 and hHSC-MLL-AF9. (**A**) An shRNA screen identifies enrichment of an shRNA targeting Trp53 (red) after treatment with IMG-7289. (**B**) GSEA was used to compare steady-state RNA-seq datasets from 3 independent untreated LSK-MLL-AF9 and 3 untreated GMP-MLL-AF9 leukemias (left and right panels). Previously published TCGA^34^ RNA-seq datasets from 27 AML patient samples with high *EVI1* expression and 27 samples with low *EVI1* expression were analyzed by GSEA (center panel). (**C**) Heatmap representation of RNA-seq expression levels of a subset of p53 target genes from 3 independent untreated LSK-MLL-AF9 and 3 untreated GMP-MLL-AF9 leukemias. (**D**) qRT-PCR analysis of Bax and Noxa mRNA upregulation in GMP-MLL-AF9, LSK-MLL-AF9 and p53^-/-^ GMP-MLL-AF9 treated for 48 hours with DMSO control or 1 μM IMG-7289. Data were normalized to DMSO control for each leukemia subtype. (**E**) Western blot analysis for p53 protein expression in GMP-MLL-AF9 and LSK-MLL-AF9 treated for 48 hours with DMSO or 1 μM IMG-7289. Histone H3 was used as loading control. (**F**) Western blot analysis for p53 in human cord blood hGMP-MLL-AF9 and hHSC-MLL-AF9 cells. Histone H3 used as loading control. (**G**) qRT-PCR analysis of p53 mRNA expression in independent GMP-MLL-AF9 and LSK-MLL-AF9 leukemias treated for 48 hours with DMSO or 1 μM IMG-7289. Data were normalized to DMSO control for each leukemia line. (**H**) p53 Western blot analysis of LSK-MLL-AF9 cells expressing control shRNA or independent Evi1 shRNAs. (**I**) NIH-3T3 cells were transiently transfected with empty vector or FLAG-tagged Evi1 plasmid constructs and treated for 3 hours with variable concentrations of doxorubicin. p53, FLAG, and histone H3 Western blots are shown. (**J**) LSK-MLL-AF9 cells were treated with increasing concentrations of the MDM2 inhibitor RG7388 for 3 days and compared to untreated GMP-MLL-AF9 to assess p53 protein abundance by Western blot. (**K**) LSK-MLL-AF9 cells were pretreated with 200 nM RG7388 and subjected to Cell Titer Glo IMG-7289 kill curve assays with GMP-MLL-AF9 and LSK-MLL-AF9 without RG7388 pre-treatment for comparison. Bar plots of IC_50_ values from 3 independent Cell Titer Glo IMG-7289 kill curve assays are shown.

We also performed RNA-seq analysis comparing the transcriptomes of murine LSK- and GMP-derived MLL-AF9 leukemias under untreated, steady-state conditions. Using gene set enrichment analysis (GSEA), we found that among the gene sets most positively enriched in LSK-derived MLL-AF9 leukemias (i.e. P53_DN.V1_UP) was one defined by p53-mutant NCI-60 cell lines compared to p53-wild type cell lines (**Figure 3b, left panel**)^31^. We also identified a gene set enriched in GMP-derived murine leukemias (**Figure 3b, right panel**) previously defined by expression of primary canonical p53 target genes^32^ (KANNAN_TP53_TARGETS_UP), which suggests that GMP-derived MLL-AF9 leukemias have higher basal transcription of canonical p53 target genes. Indeed, when represented as a heat map (**Figure 3c**), primary p53 target genes are more highly transcribed in the 3 independent GMP-derived leukemias compared to 3 independent LSK-derived leukemias. Importantly, we confirmed that these LSK-MLL-AF9 and GMP-MLL-AF9 genomes harbored wild-type p53 alleles without copy-number alterations by using targeted sequencing (**Supp. Figure 2c-d**) as well as low-pass whole genome sequencing (**Supp. Figure 2e**) approaches^33^. We next analyzed RNA-seq AML data from TCGA^34^. Out of the 179 primary AML samples that were analyzed by RNA-seq, only 4 patient samples harbored MLL gene rearrangements, and 3 of these 4 samples were *EVI1*^high^ and 1 was *EVI1*^*low*^. For this reason, we instead ranked all 179 TCGA AML patient samples in order of normalized *EVI1* expression level. From this rank list, we compared the 27 AML samples with the highest *EVI1* expression to the 27 samples with the lowest abundance of *EVI1* mRNA and applied GSEA to these cohorts. Importantly, while MLL-rearranged AML were observed in both *EVI1*^*high*^ and *EVI1*^*low*^ groups, there were a variety of other AML subtypes represented within this analysis, including but not limited to inversion(16), t(8;21), complex karyotype, NUP98-rearranged AML, t(17;19), t(3;5), and normal karyotype-AML. The same p53 gene set (i.e. P53_DN.V1_UP) as noted above had positive enrichment within *Evi1*^high^ AML samples of genes compared to *Evi1*^low^ AML samples (**Figure 3b, center panel**). These findings suggest that the transcriptional profiles of primary AML patient samples, when segregated based on *EVI1* expression level irrespective of the presence or absence of MLL gene rearrangements, recapitulate the differential p53-related gene expression profiles observed in murine MLL-AF9 leukemias based on differential cells of origin. In addition, these data suggest that the p53 mediated gene expression program may be modulated by differential *Evi1* expression.

We next assessed p53 activity with a focus on transcriptional activation of pro-apoptotic targets downstream of p53 in the setting of LSD1 inhibitor treatment. LSK- and GMP-MLL-AF9 leukemias were treated with IMG-7289 for 48 hours, after which quantitative real-time polymerase chain reaction (qRT-PCR) was performed to assess mRNA abundance of the pro-apoptotic p53 target genes, *Bax* and *Noxa* (**Figure 3d**). IMG-7289 treatment of GMP-derived, but not LSK-derived, MLL-AF9 leukemia cells resulted in significant upregulation of *Bax* and *Noxa* expression relative to control. To assess whether upregulation of these target genes was p53-dependent in GMP-derived leukemias, we generated primary GMP-derived MLL-AF9 leukemias from p53-knockout mice and found that loss of p53 in GMP-derived leukemias abrogated IMG-7289-induced upregulation of *Bax* and *Noxa*. Upregulation of p21 was seen after IMG-7289 treatment in both GMP- and LSK-derived leukemias, as well as in p53-null GMP-derived leukemias (**Supp. Figure 2f**), as p53-independent activation of p21 has been described^35,36^. These data suggest that the transcriptional output of pro-apoptotic p53 target genes is greater in GMP-derived leukemias than in LSK-derived leukemias.

We hypothesized that the increased p53 transcriptional output seen in GMP-derived MLL-AF9 leukemias could be due to increased p53 protein abundance in GMP-MLL-AF9 cells relative to LSK-MLL-AF9 cells and/or increased chromatin occupancy of p53 in GMP-MLL-AF9 cells when compared to LSK-derived leukemias. To test the first possibility, two independent GMP-MLL-AF9 and two independent LSK-MLL-AF9 cells were treated with DMSO or IMG-7289, and whole cell extracts were subjected to Western blot analysis for p53 protein abundance (**Figure 3e**). Comparison of p53 protein abundance in DMSO control lanes revealed higher basal levels of p53 protein expression in GMP-MLL-AF9 cells compared to LSK-MLL-AF9 cells. Furthermore, treatment with IMG-7289 revealed further stabilization of p53 protein in GMP-MLL-AF9 cells, which was not oberved in IMG-7289-treated LSK-MLL-AF9 cells. Extended exposure times revealed that p53 protein was indeed detectable in LSK-MLL-AF9 cells, but p53 protein stabilization remained blunted in these cells even after IMG-7289 treatment (**Supp. Figure 2g**). Similarly, Western blot analysis revealed greater p53 protein abundance in hGMP-MLL-AF9 cells (**Figure 3f**). Importantly, under the same IMG-7289 treatment conditions that caused induction of p53 protein expression in GMP-MLL-AF9 cells, no increase in p53 mRNA expression was seen in either GMP-MLL-AF9 or LSK-MLL-AF9 leukemia cells, suggesting that IMG-7289-induced p53 protein stabilization is post-transcriptionally regulated (**Figure 3g**). In fact, treatment with LSD1 inhibitor resulted in relatively fewer p53 mRNA transcripts, which may potentially be explained by selection for viable cells with less p53 activity over the course of exposure to IMG-7289.

Multiple cellular stresses can induce p53 protein stabilization via disruption of the interaction between p53 and Mdm2, the E3 ubiquitin ligase targets p53 for proteasomal degradation. The DNA damage response is one cellular stress that can induce p53 activation and stabilization. To detect differences in DNA damage response between LSK-MLL-AF9 and GMP-MLL-AF9 cells, we performed 6-hour time course during which these cells were treated with 1 μM IMG-7289 and then cell lysates were harvested for Western blot analysis to detect gamma-H2AX induction with 0, 1, 3, and 6 hours of exposure to LSD1 inhibitor (**Supp. Figure 3f**). We observed greater amounts of gamma-H2AX at 0, 1, and 3 hours post-treatment with IMG-7289 in GMP-MLL-AF9 cells relative to LSK-MLL-AF9 cells. Next, we treated NIH-3T3 cells expressing either murine Evi1 or empty vector with increasing concentrations of the proteasomal inhibitor bortezomib in order to block the degradation of ubiquitinated p53. If Evi1 were influencing p53 protein stability at the level of the proteasome or at the ubiquitination step for p53, we would expect that Evi1-overexpressing cells treated with bortezomib would have greater accumulation of mono- or poly-ubiquitinated species of p53 (**Supp. Figure 3g**). Notably, while bortezomib treatment did reveal higher molecular weight modified p53 species detectable by Western blot analysis, Evi1 overexpression in NIH-3T3 cells did not alter the abundance of these species when compared to empty vector-transduced cells. These data suggest that, while the DNA damage response upstream of the p53 pathway may be influenced by cell of origin, Evi1 overexpression did not alter proteasome-dependent stability of p53, suggesting Evi1 modulates p53 stability via a proteasome-independent mechanism.

Analysis of the TCGA AML samples (**Figure 3b, center panel**), together with the negative correlation between *Evi1* expression level and p53 protein abundance, led us to hypothesize that *Evi1* functions as an upstream modulator of p53 protein stability. Knockdown of *Evi1* in LSK-MLL-AF9 cells resulted in increased p53 protein expression relative to non-targeting shRNA control (**Figure 3h**). Moreover, NIH-3T3 cells expressing FLAG-Evi1 exhibited less p53 protein stabilization after treatment with doxorubicin when compared to cells transfected with empty vector (**Figure 3i**). We also treated LSK-MLL-AF9 cells with the MDM2 inhibitor RG7388 to induce p53 protein stabilization at levels comparable to GMP-MLL-AF9 cells (**Figure 3j**). Treatment with RG7388 was sufficient to restore LSD1 inhibitor sensitivity to LSK-MLL-AF9 cells (**Figure 3k**). These experiments demonstrate that cell of origin of leukemic transformation influences p53 protein stability in an Evi1-dependent manner.

### Differential sensitivity of MLL-AF9 leukemias to LSD1 inhibition is determined by p53

To determine if the abundance of p53 protein in LSK- and GMP-derived leukemias contributes to differential sensitivity to LSD1 inhibition, we knocked down p53 in GMP-MLL-AF9 cells and assessed for sensitivity to IMG-7289 treatment. When compared to non-targeting shRNA control, p53 knockdown in GMP-MLL-AF9 leukemias significantly enhanced resistance to IMG-7289 (**Figure 4a**). We then isolated normal GMPs from p53-knockout mice and used these cells to generate MLL-AF9 primary leukemias. When compared to p53-wild type GMP-derived leukemias, p53-null GMP-derived leukemias were more resistant to IMG-7289 (**Figure 4b**). Knockdown with p53 shRNA – but not Renilla shRNA – in GMP-MLL-AF9 cells abrogated IMG-7289-induced mitochondrial depolarization assessed by TMRE staining (**Figure 4c-d**) as well as apoptosis induction by IMG-7289 treatment measured by Annexin V staining (**Figure 4e**).

**Figure 4.**
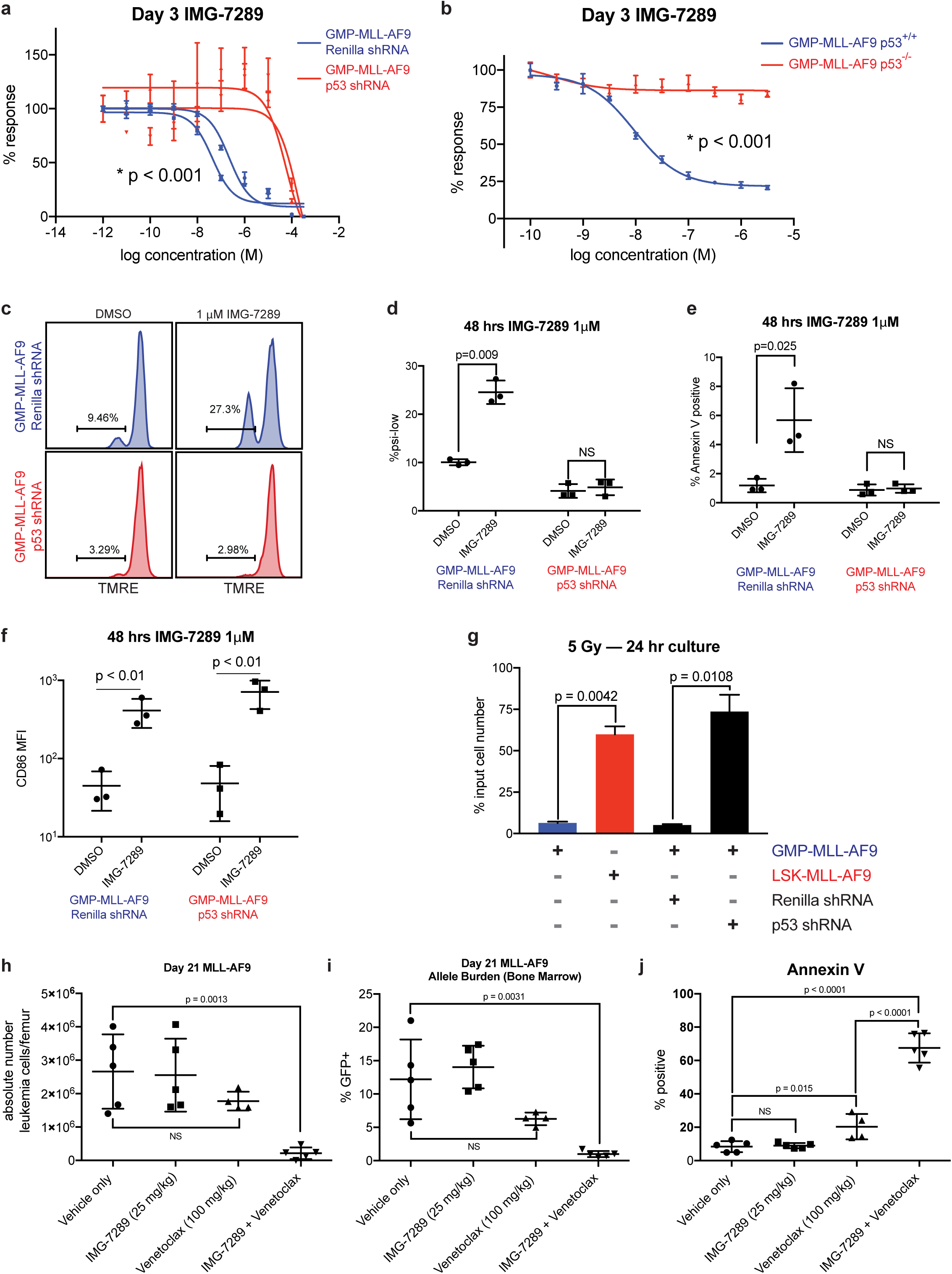
p53 loss in sensitive GMP-MLL-AF9 leukemias induces resistance to the pro-apoptotic but not pro-differentiation effects of LSD1 inhibition. (**A**) GMP-MLL-AF9 cells were transduced with shRNAs targeting Renilla luciferase control or p53 and assayed in Cell Titer Glo IMG-7289 kill curves. Data shown are representative of 4 independent experiments. (**B**) Primary MLL-AF9 leukemias arising from GMPs derived from either p53 wild-type or p53^-/-^ mice were assessed for IMG-7289 sensitivity in vitro by Cell Titer Glo kill curves. Data shown are representative of 4 independent experiments. (**C**) GMP-MLL-AF9 cells expressing shRNAs targeting Renilla or p53 were treated for 48 hours at 1 μM IMG-7289 and TMRE fluorescence was measured by flow cytometry. Statistical analyses summarized in (**D**). Data shown are representative of 4 independent experiments. (**D**) Summary scatter plots depicting the percentage of cells within the Ψ^low^ gate defined by TMRE flow cytometry in C. (**E**) Summary plots showing the proportion of Annexin V^+^ apoptotic cells after GMP-MLL-AF9 cells expressing shRNAs targeting Renilla or p53 were treated for 48 hours at 1 μM IMG-7289. (**F**) GMP-MLL-AF9 cells expressing shRNAs targeting Renilla or p53 were treated for 48 hours at 1 μM IMG-7289 and assessed for cell surface CD86 expression by flow cytometry. Scatter plots depicting mean fluorescence intensity of CD86 staining is shown. Data shown are representative of 3 independent experiments. (**G**) 2×10^5^ parental GMP-MLL-AF9, GMP-MLL-AF9 expressing Renilla shRNA, GMP-MLL-AF9 expressing p53 shRNA, and LSK-MLL-AF9 cells were irradiated (5 Gy). Viable cells were enumerated after culturing for 24 hours post-irradiation. Bar plots represent viable cell counts as a percentage of input cell number. (**H**) C57BL/6 wild-type mice were engrafted with resistant LSK-MLL-AF9 cells and treated via oral gavage with vehicle (N=5), IMG-2789 alone (25 mg/kg, N=5), venetoclax alone (100 mg/kg, N=5), or combination IMG-2789 (25 mg/kg) and venetoclax (100 mg/kg, N=5) during a 14-day treatment window. At day 21 post-transplant, bone marrow cells were harvested and viable GFP^+^ leukemia cells were enumerated per femur. (**I**) Percentages of GFP^+^ AML cells within the bone marrow are shown, and (**J**) proportions of Annexin V^+^GFP^+^ apoptotic leukemia cells are shown.

One well-characterized consequence of LSD1 inhibition is induction of myeloid maturation or differentiation, which is associated with cell surface upregulation of the type I transmembrane protein, CD86^11,37-40^. To determine if perturbation of the p53 pathway affects LSD1 inhibitor-mediated upregulation of cell surface CD86 expression, we treated GMP-MLL-AF9 cells expressing shRNAs targeting either p53 or Renilla with IMG-7289 and measured cell surface induction of CD86. Knockdown of p53 in GMP-MLL-AF9 cells did not affect IMG-7289-induced CD86 upregulation measured by mean fluorescence intensity of CD86 staining (**Figure 4f**). These data suggest that perturbation of p53 expression in GMP-MLL-AF9 cells is sufficient to induce resistance to the pro-apoptotic effects of IMG-7289 treatment but is insufficient to reverse the pro-differentiation effects of LSD1 inhibition, thereby functionally separating apoptosis induction and differentiation in response to LSD1 inhibition.

We hypothesized that the differential p53 activity seen in leukemias arising from distinct cells of origin would also result in differential sensitivity to other stimuli that trigger p53 activation, as has been shown for cytotoxic chemotherapy such as doxorubicin^3-5^. GMP-MLL-AF9 and LSK-MLL-AF9 leukemia cells were exposed to ionizing radiation at a dose of 5 Gy and cultured for 24 hours post-irradiation, and viable cells were harvested and enumerated. LSK-MLL-AF9 leukemias were more resistant to ionizing radiation, with a greater number of viable cells counted post-irradiation when compared to GMP-MLL-AF9 leukemias (**Figure 4g**). Furthermore, this effect was p53-dependent, as p53 knockdown induced resistance to irradiation in GMP-derived leukemias. Taken together, these findings demonstrate that the pro-apoptotic effects of LSD1 inhibition, similar to other more classical genotoxic stresses (e.g. chemotherapy, ionizing radiation) are p53-dependent.

The observation that p53 activity can be modulated by Evi1 is also consistent with clinical data ascribing chemoresistance, higher rates of relapse, and poorer overall prognosis to AML exhibiting high *EVI1* expression^7,41,42^. We hypothesized that the addition of venetoclax, a BH3 mimetic and small-molecule inhibitor of Bcl-2, to LSD1 inhibition would enhance anti-leukemic activity and apoptotic priming in LSK-derived MLL-AF9 leukemias *in vivo* and reverse resistance to LSD1 inhibition. Venetoclax is now approved for use in combination with hypomethylating agents for AML patients who were deemed to be unfit for standard induction in the upfront setting with very promising response rates even among AML patients with adverse-risk molecular features^43-45^. We postulated that targeting Bcl-2 in combination with LSD1 inhibition would enhance apoptosis induction in LSK-derived leukemias that exhibit blunted p53 activity and apoptotic priming. To test this hypothesis, we performed *in vitro* growth inhibitory assays comparing LSK- and GMP-derived leukemias in a two-drug matrix using variable concentrations of IMG-7289 and venetoclax (**Supp. Figure 3e**). From these data, we derived synergy scores using Bliss independence models (**Supp. Table 2**) and found that the lowest concentrations of IMG-7289 and venetoclax at which synergy (defined as Bliss score > 10) was achieved were 1 uM and 1 nM, respectively. This synergy was seen for LSK-MLL-AF9 as well as for GMP-MLL-AF9 leukemias. To test this synergy *in vivo*, we transplanted recipient mice with resistant LSK-MLL-AF9 cells and began treating cohorts via oral gavage with vehicle, IMG-7289, venetoclax, or combination IMG-7289 and venetoclax therapy for 14 days. While administration of IMG-7289 or venetoclax as monotherapy did not result in a statistically significant reduction in the absolute numbers of GFP^+^ LSK-MLL-AF9 cells per femur in each recipient mouse, combination IMG-7289 and venetoclax therapy reduced absolute leukemic blasts per femur (**Figure 4h**). Similarly, combination therapy induced a significant reduction in the percentage of GFP^+^ leukemic blasts in the bone marrow (**Figure 4i**). Furthermore, venetoclax monotherapy enhanced the proportion of Annexin V^+^ MLL-AF9 cells, but the addition of IMG-7289 to venetoclax further enhanced the fraction of apoptotic leukemic blasts within the bone marrow compartment (**Figure 4j**). These data suggest that the addition of venetoclax to IMG-7289 can enhance anti-leukemic activity in *Evi1*^high^ LSK-derived MLL-AF9 leukemia cells.

## Discussion

In this study, we show that leukemias initiated by MLL-AF9 arising from distinct mouse or human hematopoietic stem/progenitor compartments exhibit markedly different therapeutic susceptibilities to cytotoxic chemotherapy, ionizing radiation, and LSD1 inhibition. These cell-of-origin-dependent effects are conferred by differential p53 protein levels, which is related to differential *Evi1* expression in murine and human leukemic contexts. Our data suggest that cell of origin can influence p53 activity – not through genomic or transcriptional aberrations in p53 – but rather through regulating p53 protein abundance. While it currently remains unclear how p53 protein abundance is controlled, a number of well-characterized post-translational modifications, including but not limited to MDM2-mediated ubiquitination of p53, could be mediating this effect and well suited to subsequent functional interrogation.

Previous studies have demonstrated leukemogenic fusion genes, such as MLL-AF9, have the capacity to induce transformation of murine hematopoietic stem cells and of myeloid progenitors, and that these leukemias have many similarities in their biology and in the immunophenotype of the cells capable of propagating the disease^4,46^. However, our data suggest that specific features of the disease, including epigenetic regulation of the *Evi1* locus and sensitivity to LSD1 inhibition or to genotoxic stress, is governed by the cell type which acquires the leukemia-defining genetic alteration. Notably, cell of origin does not impact the responsiveness of AML cells to epigenetic therapies which induce therapeutic efficacy primarily through induction of differentiation, such as Dot1L inhibition, and does not impact the ability of LSD1 inhibition to promote myeloid maturation of leukemic cells based on cell of origin. Furthermore, our studies of umbilical cord blood-derived MLL-AF9 leukemias established definitively that (1) both human HSCs and GMPs can be transformed by MLL-AF9 and that (2) human MLL-AF9 leukemias arising from different cells of origin, like murine leukemias, exhibit differential apoptotic priming and p53 protein stability. Subsequent studies will delineate whether cell-of-origin impacts the response to additional anti-leukemic therapeutic modalities, and if cell of origin modulates therapeutic dependencies in other AML subtypes with different somatic alterations.

Our functional studies led us to evaluate whether pro-apoptotic BH3 mimetics such as venetoclax, in combination with LSD1 inhibition, could overcome the drug resistance seen in *Evi1*^high^ leukemias. Taken together, our findings delineate a role for *Evi1* in perturbing p53 protein stability in an otherwise p53 genetically wild-type malignancy, which influences sensitivity to chemotherapy and targeted epigenetic therapies. Our work identifies a particular pathway of drug resistance conferred through *Evi1* that can be circumvented by therapeutic combinations that short-circuit apoptotic priming through Bcl-2.

## Methods

Methods and any associated references are available in the online version of the paper.

## Acknowledgements

We thank members of the Levine, Armstrong, and Lowe laboratories for discussions. We thank Yanming Zhang for assistance with cytogenetic analysis of AML patient samples. S.F.C. is supported by a Scholar Award from the American Society of Hematology, a Momentum Fellowship Award from The Mark Foundation for Cancer Research, a Young Investigator Award from The Truth 365 and the Rally Foundation for Childhood Cancer Research, a Young Investigator Award from the Conquer Cancer Foundation, a Career Development Program Fellow Award (5453-17) from the Leukemia & Lymphoma Society, and a Memorial Sloan Kettering Cancer Center Clinical Scholars Biomedical Research Fellowship. R.L.L. is supported by National Cancer Institute grants P01 CA108671 and R35197594 as well as a Leukemia & Lymphoma Society Specialized Center of Research grant. S.A.A. is supported by grants from the National Institutes of Health (NIH), National Cancer Institute grants PO1 CA66996 and R01 CA140575, and Gabrielle’s Angel Research Foundation. Studies supported by Memorial Sloan Kettering core facilities were supported in part by Memorial Sloan Kettering Cancer Center Support Grant/Core Grant P30 CA008748.

## Author contributions

S.F.C., H.Y.R., A.L., R.L.L., and S.A.A. conceived the project, designed the experiments, analysed the data, and wrote the manuscript. S.F.C, S.H.C., S.P., M.C., C-H.H., and C-W.C. performed the experiments with technical assistance from K.B.C., V.D., M.K., and M.R. ChIP-seq and RNA-seq computational analyses were performed by R.P.K. and L.Z. J.Z., T.B., and S.W.L. generated custom shRNA libraries and performed copy number analyses. A.D.G., J.L.G., E.M.S., M.S.T., and C.F. assisted with management of clinical data and specimens. F.S. assisted with pathological assessment of biospecimens.

## Supplemental Figure Legends

**Supplemental Figure 1.**

(**A**) Independent murine LSK- and GMP-derived MLL-AF9 leukemias were treated for 3 days with variable concentrations of doxorubicin *in vitro*. Cell Titer Glo assays were used to quantitate viable cells cultured in triplicate wells and percent response was derived by normalizing to corresponding cells cultured in DMSO vehicle control. (**B**) Independent murine LSK- and GMP-derived MLL-AF9 leukemias were passaged and absolute cells were enumerated every 3 days over a 12-day period, and (**C**) doubling times were derived from this culture period. (**D**) LSK- and GMP-derived MLL-AF9 leukemia cells were subjected to IMG-7289 kill curve assays after culture over time spanning a 15-day treatment period. Bar plots represent IC_50_ values derived from independent MLL-AF9 lines. (**E**) Kill curve assays assessing LSK- and GMP-MLL-AF9 cells after 3 days of treatment with the LSD1 inhibitor IMG-98. (**F**) Kill curve assays assessing LSK- and GMP-MLL-AF9 cells after 15 days of treatment with the DOT1L inhibitor EPZ-5676. (**G**) Representative flow plots from human cord blood HSC- and GMP-derived MLL-AF9 leukemia cells harvested from the bone marrow of engrafted NSG recipient mice, showing expression of the myeloid marker CD33 after gating on GFP^+^ leukemic blasts.

**Supplemental Figure 2.**

(**A**) Summary bar plots depicting comprehensive BH3 profiling data spanning all peptides and compounds tested. Apoptosis induction in independent GMP-MLL-AF9 and LSK-MLL-AF9 murine AML lines was assessed by cytochrome C release. (**B**) Summary bar plots depicting BH3 profiling of human cord blood HSC- and GMP-MLL-AF9 cells. (**C**) Bar plots showing sequencing coverage spanning all 11 exons of the Trp53 gene when independent GMP-MLL-AF9 and LSK-MLL-AF9 cells are subjected to (**D**) next-generation targeted sequencing analysis of a panel of 572 murine cancer-associated genes. Mutations called are shown with variant allele fractions (VAFs) shown within each box corresponding to the specific MLL-AF9 cell line (top). (**E**) Low-pass whole genome sequencing was performed on independent GMP-MLL-AF9 and LSK-MLL-AF9 lines. Zoomed-in-view of the copy number of independent LSK-MLL-AF9 (red) and GMP-MLL-AF9 (blue) leukemias at the 11B3 locus (syntenic to human 17q13.1) showing retention of both alleles of the Trp53 gene (vertical gray bar). (**F**) qRT-PCR analysis of p21 mRNA upregulation in GMP-MLL-AF9, LSK-MLL-AF9 and p53^-/-^ GMP-MLL-AF9 treated for 48 hours with DMSO control or 1 μM IMG-7289. Data were normalized to DMSO control for each leukemia subtype. (**G**) Prolonged exposure of p53 Western blot shown in **Figure 3E**.

**Supplemental Figure 3.**

(**A**) Western blot analysis showing Bcl-2 and Histone H3 (loading control) protein expression in 3 independent GMP-MLL-AF9 cell lines transduced with either empty vector or a Bcl-2 expression vector after selection with puromycin. (**B**) BH3 profiling of murine GMP-derived MLL-AF9 leukemia cells expressing empty vector (black) or Bcl-2 (red). (**C**) Scatter plots depicting IC_50_ values derived from independent Cell Titer Glo IMG-7289 kill curve assays using GMP-MLL-AF9 cells expressing empty vector or Bcl-2. (**D**) Bar plots depicting normalized viable cell counts 24 hours after exposure to 10 nM doxorubicin, comparing GMP-MLL-AF9 cells transduced with either empty vector (purple bar) or Bcl-2 (red bar). Data shown are representative of 4 independent experiments. (**E**) Viable cell counts (normalized to DMSO control) represented in heat map form for LSK-MLL-AF9 (left) and GMP-MLL-AF9 (right) treated for 72 hours with variable concentrations of venetoclax (y-axis) and IMG-7289 (x-axis). Data are representative of 3 indepdendent experiments. (**F**) Western blot analysis showing gamma-H2AX and histone H3 (loading control) protein expression in GMP-MLL-AF9 (indicated by red bars) and LSK-MLL-AF9 (indicated by blue bars) cells treated over a 6-hour time course with 1 μM IMG-7289. Data are representative of 3 indepdendent experiments. (**G**) Western blot analysis showing p53 and histone H3 (loading control) protein expression in NIH-3T3 cells transduced with either empty vector or a codon-optimized version of murine Evi1 and treated for 24 hrs with increasing concentrations of bortezomib (0, 0.1, 1, and 10 μM). Data are representative of 3 indepdendent experiments.

## Online Methods

### Mice

Laboratory mice were housed at the Memorial Sloan Kettering Cancer Center (MSKCC) animal facility. All animal procedures were approved by the Institutional Animal Care and Use Committee. C57BL/6 mice (CD45.2, Charles River) were used as transplantation recipients. p53 knockout mice (002101) were obtained from The Jackson Laboratory. Both female and male mice were used in our leukemia studies and no obvious sex-dependent differences were observed.

### Human cord blood

Normal cord blood units, designated for research use, were obtained from the New York Blood Center (New York, NY) under IRB approval from the National Cord Blood Program. Both male and female samples were randomly assigned and used in this study. The experiments were approved by the institutional biosafety committees at MSKCC.

### Cell lines and cell culture

293T and NIH-3T3 cell lines were maintained in DMEM medium supplemented with 10% fetal bovine serum (FBS) and 1% penicillin-streptomycin. Cells were maintained in a humidified incubator at 37° C, 5% CO2.

### Plasmid constructs

A codon-optimized, doxycycline-inducible version of murine Evi1 (pRRL.PPT.Tet.Evi1.IRES.EGFP.pre) and empty vector control plasmid (pRRL.PPT.Tet.Evi1.IRES.EGFP.pre) were kind gifts from Dr. Olga Kustikova, Dr. Christopher Baum, and Dr. Axel Shamback (Medizinische Hochschule Hannover, Hannover, Germany). Generation of these reagents were previously described^47^.

### Murine and human MLL-AF9 leukemia models

Murine bone marrow cells were obtained from femurs, tibias, hips, and spines and stained with a biotinylated lineage antibody cocktail (559971, BD Biosciences), Sca1-PE-Cy7, cKit-APC, and F_c_Rg-Pacific Blue antibodies. The immunophenotypes of LSK and GMP cells are lineage^-^Sca1^+^cKit^+^ and lineage^-^cKit^+^Sca1^-^ CD34^+^F_c_Rg^+^, respectively. LSK and GMP cells were isolated using a FACS Aria (BD Biosciences) and cultured for 24 hours in IMDM medium containing 15% FBS, 1% penicillin-streptomycin, 2 mM L-glutamine, 20 ng/mL rm-SCF, 10 ng/mL rmIL-3, 10 ng/ml rmIL-6. Spinfections were then performed by adding retrovirus carrying MLL-AF9-ires-GFP in the media containing 8 μg/mL polybrene to either LSK or GMP cells and centrifuging at 1,400 x g for 90 minutes. MLL-AF9 immortalized LSK or GMP cells were expanded *in vitro* and a minimum of 2×10^5^ GFP^+^ cells were transplanted into lethally irradiated (950 cGy) syngeneic recipient C57BL/6 mice along with 2×10^5^ bone marrow support cells from wild-type mice. Murine AML cells were harvested from bone marrow of moribund mice and maintained in culture. Human cord blood CD34^+^ cells were enriched by microbeads (Miltenyi Biotec). The immunophenotypes of hHSC and hGMP cells are lineage^-^CD34^+^CD38^-^ and lineage^-^CD34^+^CD38^+^CD45RA^+^CD110^−^, respectively. hHSC and hGMP cells were sorted and cultured in IMDM medium containing 20% FBS, 1% penicillin-streptomycin, 2 mM L-glutamine, β-mercaptoethanol, 20 ng/mL each of rh-IL-6, rhTPO, rhSCF, rhGM-CSF, and 10 ng/mL IL-3. After 24 hours in culture, hHSC and hGMP cells were retrovirally transduced with MLL-AF9-ires-GFP as was done to generate murine leukemias. Immortalized cells were expanded *in vitro*. To generate fully transformed leukemia cells, a minimum of 1.3×10^5^ GFP^+^ MLL-AF9-expressing cells were transplanted into sublethally irradiated (250 cGy) NOD/SCID/IL-2Rg_c_-null (NSG) mice. AML cells were harvested from bone marrow of moribund mice.

### Chromatin immunoprecipitation

For ChIP-seq experiments, murine AML or primary human AML patient samples were fixed with 1% formaldehyde for 10 minutes and quenched with 0.125 M glycine for 5 minutes. Fixed cells were washed twice with ice-cold phosphate-buffered saline (PBS). Washed cells were then resuspended in ChIP lysis buffer and sheared using an E220 focused-ultrasonicator (Covaris). Appropriate amounts of antibodies were added to sheared chromatin and incubated overnight at 4° C. Immune complexes were collected with protein A/G agarose and washed sequentially in low-salt wash buffer (20 mM Tris pH 8.0, 150 mM NaCl, 0.1% SDS, 1% Triton X-100, 2mM EDTA), high-salt wash buffer (20 mM Tris pH 8.0, 500 mM NaCl, 0.1% SDS, 1% Triton X-100, 2mM EDTA), LiCl wash buffer (10 mM Tris pH 8.0, 250 mM LiCl, 1% NP-40, 1% sodium deoxycholate, 1 mM EDTA), and TE. Chromatin was eluted in elution buffer (1% SDS, 0.1 M NaHCO3), and then reverse cross-linked with 0.2 M NaCl at 65° C for 4 hours. DNA was purified with a PCR purification kit (QIAGEN). Libraries were prepared using a ThruPLEX DNA-seq Kit (Rubicon Genomics). The DNA library was validated using a TapeStation (Agilent Technologies) and was quantified using a Qubit 2.0 Flurometer (Thermo Fisher Scientific). Libraries were pooled and sequenced on an Illumina HiSeq2000.

### RNA isolation and analysis

Total RNA was extracted from murine AML or primary human AML patient samples using the RNeasy Mini kit (QIAGEN). cDNA was synthesized using the Tetro cDNA synthesis kit (Bioline). cDNA fragments were quantified by a TaqMan Gene expression assay (Applied Biosystems) or by SYBR Green Real-time PCR with a ViiA 7 Real-time PCR system. RNA-seq libraries were prepared by using a ThruPLEX DNA-seq Kit (Rubicon Genomics). The DNA library was validated using TapeStation (Agilent Tehnologies) and was quantified using a Qubit 2.0 Fluorometer (Thermo Fisher Scientific). Libraries were pooled and sequenced on Illumina HiSeq2000.

### shRNA screen

A customized shRNA library (TRMPV-Neo system) focused on mouse chromatin-regulating genes, together with control shRNAs. After sequence verification, the virus library containing 2,252 shRNAs targeting 468 genes (4 to 6 shRNAs per gene) was pooled. The virus pool was transduced into monoclonal mouse MLL-AF9 (MA9) leukemic cells, stably expressing rtTA3 (Tet-on system), at a viral titer that on average causes a single viral transduction per cell and at which each shRNA is represented in at least 2,000 cells. The infected cells were selected for 2 days in 1 mg/mL neomycin (G418 sulfate, Corning), and subsequently, shRNA expression was induced by adding 1 μg/mL doxycycline (Sigma-Aldrich). The shRNA-expressing cells (dsRed and Venus double positive) were sorted (T0) using a FACS ARIA (BD) and cultured for 6 days (T6) in either DMSO or 1 μM IMG-7289. The integrated shRNA sequences in T0 and T6 cell samples were assessed by high-throughput sequencing (HiSeq) using the Illumina Next-Gen Sequencing HiSeq platform (Illumina).

### Computational analysis

ChIP-seq reads were trimmed for quality and Illumina adapter sequences with ‘trim_galore’ then aligned to either human assembly hg19 or mouse assembly mm9 with bowtie2 using the default parameters. The tool MarkDuplicates (http://broadinstitute.github.io/picard/) was used to remove ChIP-seq reads with the same start site and orientation. RNA-seq reads were similarly trimmed and then aligned to hg19 or mm9 with STAR using the default parameters. Genome browser profiles were created by extending each read to the average library fragment size and then computing density using the BEDTools suite (http://bedtools.readthedocs.io). Raw count tables for all transcripts were created using HTSeq v0.6.1. Expression dynamics were evaluated with DESeq2 using default library size factor normalization. The resulting log2 fold changes were then used as input for GSEA using the ‘preranked’ option.

### Quantification and statistical analyses

Error bars in all data shown represent standard deviation. Determination of statistical significance was calculated using a Student’s t test or one-way ANOVA using Prism 7 software (GraphPad).

