## Supplementary figures and images for "Leukemia cell of origin influences apoptotic priming and sensitivity to LSD1 inhibition"

### Supplemental Figures

#### Supplemental Figure 1

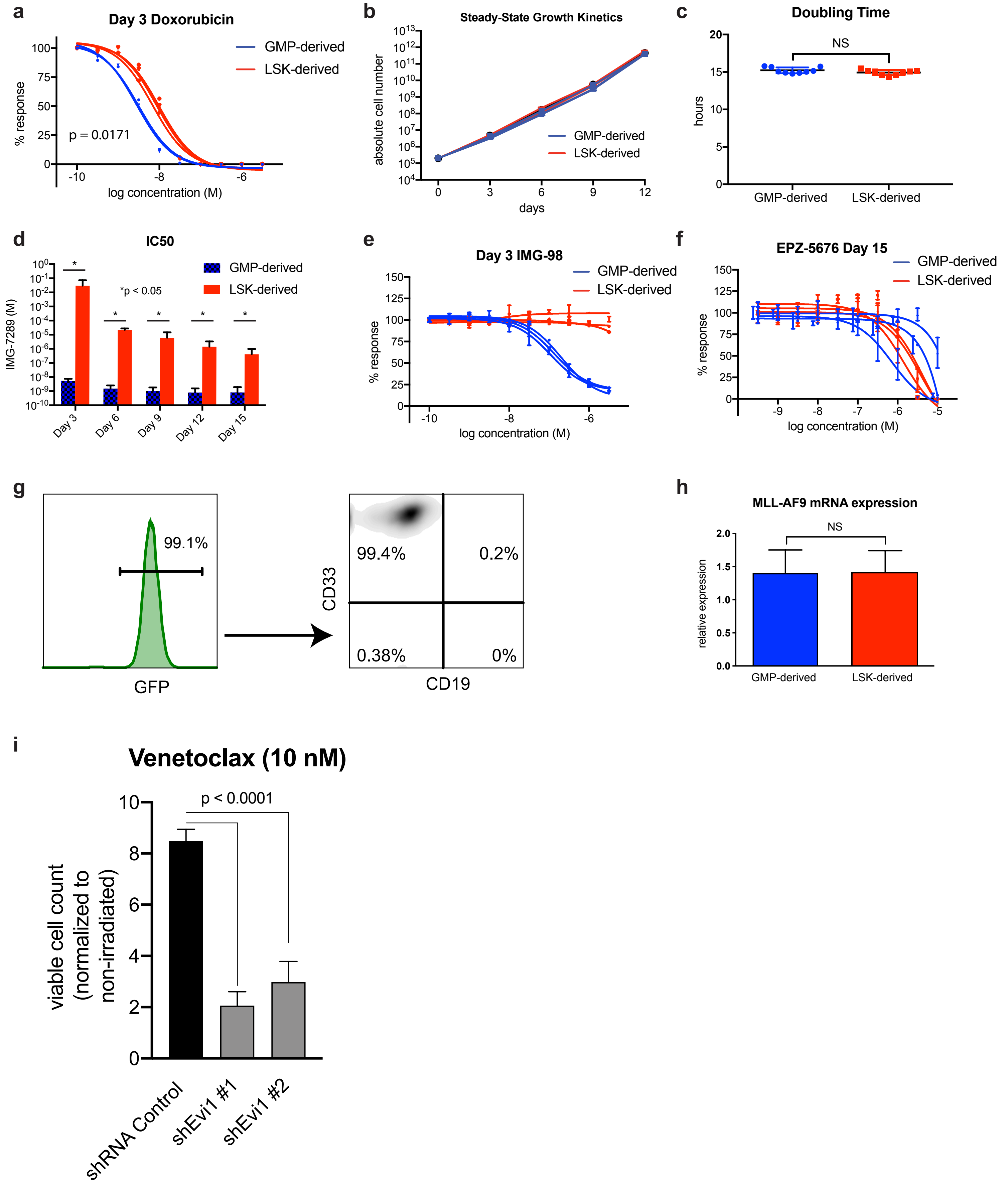

#### Supplemental Figure 2

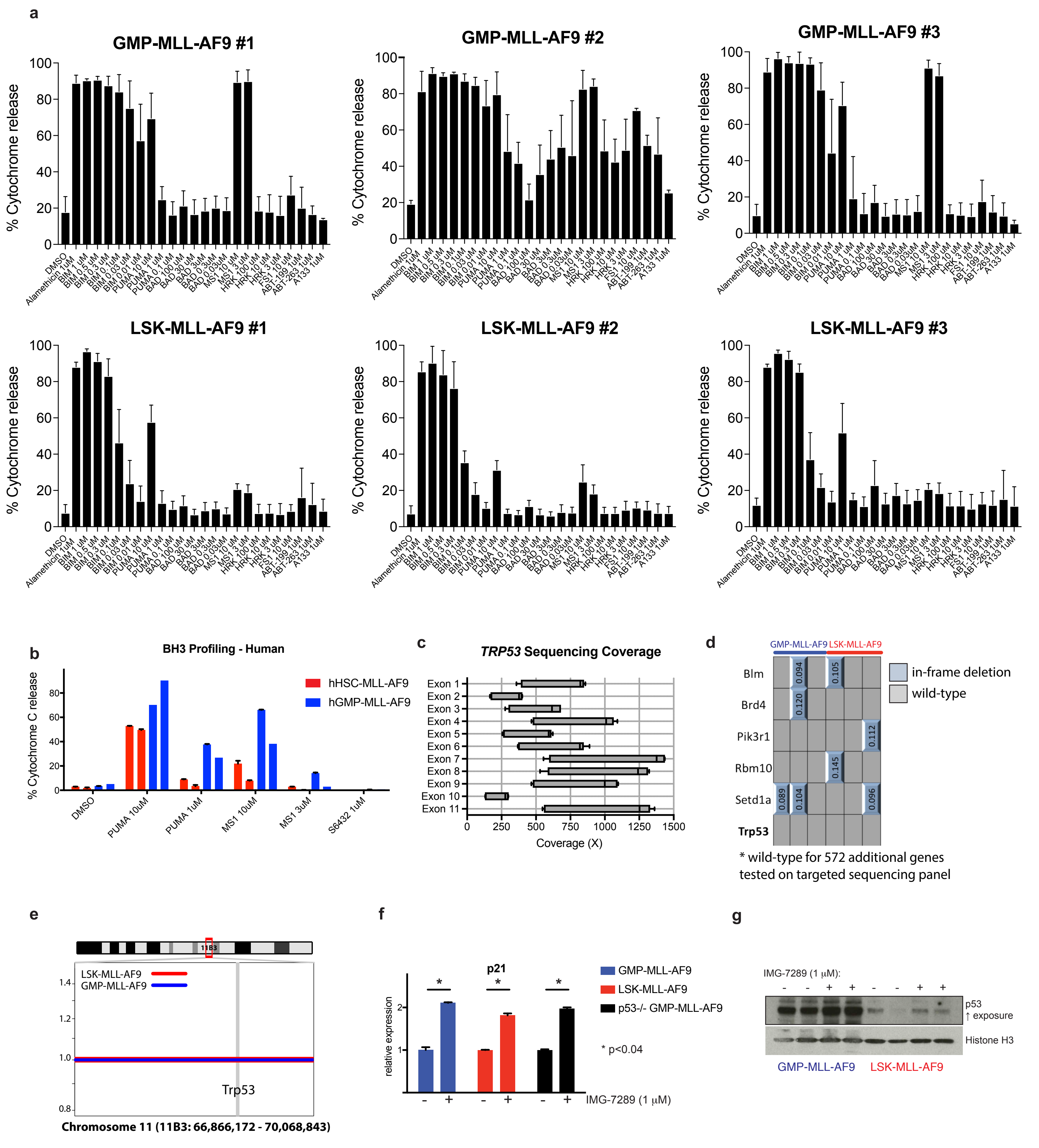

#### Supplemental Figure 3

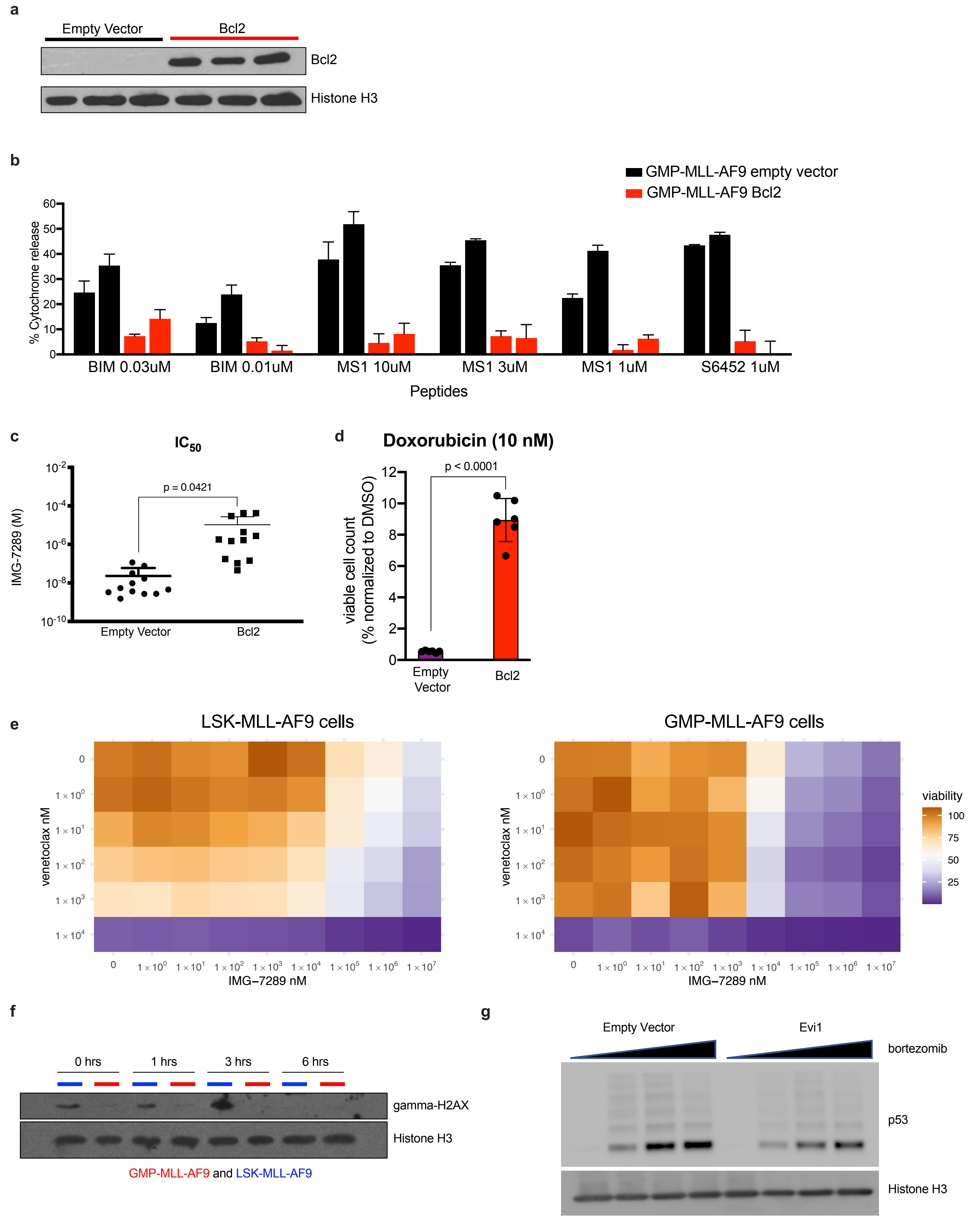
